# A Sox17 downstream gene *Rasip1* is involved in the hematopoietic activity of intra-aortic hematopoietic clusters in the midgestation mouse embryo

**DOI:** 10.1101/2022.11.01.513960

**Authors:** Gerel Melig, Ikuo Nobuhisa, Kiyoka Saito, Ryota Tsukahara, Ayumi Itabashi, Yoshiakira Kanai, Masami Kanai-Azuma, Mitsujiro Osawa, Motohiko Oshima, Atsushi Iwama, Tetsuya Taga

## Abstract

**Background:** During mouse embryonic development, definitive hematopoiesis is first detected around embryonic day (E) 10.5 in the aorta-gonad-mesonephros (AGM) region. Hematopoietic stem cells (HSCs) arise in the dorsal aorta’s intra-aortic hematopoietic cell clusters (IAHCs). We have previously reported that a transcription factor Sox17, is expressed in IAHCs, and that, among them, CD45^low^c-Kit^high^ cells have high hematopoietic activity. Furthermore, forced expression of Sox17 in this population of cells can maintain the formation of hematopoietic cell clusters. However, how Sox17 does so, particularly downstream signaling involved, remains poorly understood. The purpose of this study is to search for new Sox17 targets which contribute to cluster formation with hematopoietic activity.

**Methods:** RNA-sequencing (RNA-seq) analysis was done to identify genes that are up- regulated in Sox17-expressing IAHCs as compared with Sox17-negative ones. Among the top 7 highly expressed genes, Rasip1 which had been reported to be a vascular-specific regulator was focused on in this study and firstly the whole-mount immunostaining was done. We conducted a luciferase reporter assay to identify the Sox17 binding site for Rasip1 gene induction. We also analyzed the cluster formation and the multi-lineage colony forming ability of Rasip1-transduced cells and Rasip1-knockdown Sox17-transduced cells.

**Results:** The increase of the *Rasip1* expression level was observed in Sox17-positive CD45^low^c-Kit^high^ cells as compared with the Sox17-nonexpressing control. Also, the expression level of the *Rasip1* gene was increased by the Sox17-nuclear translocation. Rasip1 was expressed on the membrane of IAHCs, overlapping with the endothelial cell marker, CD31, and hematopoietic stem/progenitor marker (HSPC), c-Kit. Overexpression of Rasip1 in CD45^low^c-Kit^high^ cells led to a significant but transient increase in hematopoietic activity, while Rasip1 knock-down in Sox17-transduced cells decreased the cluster formation and diminished the colony-forming ability.

**Conclusions:** Rasip1 knockdown in Sox17-transduced CD45^low^c-Kit^high^ cells displayed a significant decrease in the multi-lineage colony forming ability and the cluster size. Rasip1 overexpression in Sox17-untransduced CD45^low^c-Kit^high^ cells led to a significant but transient increase in the multi-lineage colony forming ability, suggesting the presence of a cooperating factor for sustained hematopoietic activity.

## Background

During mouse ontogeny, fetal hematopoiesis is initiated from blood islands of the yolk sac that wraps around embryos [1], which is termed primitive hematopoiesis [2]. Definitive hematopoiesis is first detected around embryonic day (E) 10.5 in the aorta-gonad-mesonephros (AGM) region [3]. HSC-containing intra-aortic hematopoietic cell clusters (IAHCs) are found attached to the inside wall of the dorsal aorta [3] and arise from a specialized subset of the endothelium, the so-called hemogenic endothelium [4], which has the differentiation capacity to produce endothelial cells and hematopoietic cells [5]. Cells in IAHCs express various marker proteins. Among them, endothelial cell markers such as vascular endothelial-cadherin (VE-cad) and CD31, are expressed on cells located on the vessel wall side of IAHCs. A hematopoietic cell marker CD45 is expressed on cells of the vessel lumen side of IAHCs, while a hematopoietic stem/progenitor cell (HSPC) marker c-Kit is expressed on cluster cells [5-8]. Mice deficient in the Runx-1 transcriptional factor, which is essential for definitive hematopoiesis, show no such cell clusters in the dorsal aorta at midgestation of mouse embryo [5, 8, 9]. These expression patterns of marker proteins in IAHCs imply that specification of the hematopoietic lineage occurs within the cell cluster [10].

The expression of CD45 is generally recognized as a marker of all hematopoietic cells except for mature erythrocytes and platelets. CD45 is used, in some cases together with a hematopoietic stem/progenitor cell (HSPC) marker c-Kit, to fractionate hematopoietic stem/progenitor cells (HSPCs) in the AGM region at midgestation. In previous studies, populations with negative and low-level CD45-expression have long-term repopulating activity [6], whereas the CD45-expressing population has differentiation ability [11, 12]. Our previous studies [13, 14] showed that the cells in the primary culture of dissociated AGM regions [15], which appears to be an in vitro reproduction of hematopoiesis in the AGM region, can be separated into at least CD45^low^c-Kit^+^, CD45^low^c-Kit^−^, and CD45^high^c-Kit^low/-^ populations based on the expression profile of CD45 and c-Kit [13, 14]. The CD45^low^c-Kit^high^ cells have the highest hematopoietic capacity in vitro and the long-term reconstituting ability in vivo [16]. Then, the hematopoiesis in the AGM region rapidly decreases later than E11.5 and disappeared at E13.5, and the main site of hematopoiesis shifts to the fetal liver at E12.5 [16, 17].

An endodermal transcription factor, Sry-related high-mobility-group (HMG) box 17 (Sox17), also the other F-group (SoxF) proteins, Sox7 and Sox18, were expressed in E10.5 IAHCs [10]. Sox17, which a marker of endodermal cells and a transcriptional regulator containing a DNA binding domain called the HMG box [18, 19]. Sox17-deficient embryos have a defect in tube formation of the endodermal-derived gut and are lethal at midgestation [20]. Sox17 is expressed in cells occupying the vessel wall side of the hematopoietic cell cluster in the dorsal aorta, which is specifically expressed in endothelial cells [10, 21]. Also, mouse embryonic stem (ES) cell analysis showed that enforced Sox17-expressing cells in embryonic bodies (EBs) differentiate into hemogenic endothelial cells and hematopoietic cells [10]. Conditional knockout mice lacking Sox17 demonstrated that Sox17 is important for fetal hematopoiesis, but not adult hematopoiesis [16]. Sox17^+^VE^+^ cells from the E11.5 AGM region showed multilineage hematopoietic reconstitution in mice [21]. These expression patterns imply that Sox17 regulates the development of the early stages of HSPCs.

We have previously reported that the CD45^low^c-Kit^high^ cell population in the E10.5 AGM region although minor in size but displays the highest hematopoietic activity among all the populations classified by CD45 and c-Kit expression levels [16]. Forced expression of Sox17 in E10.5 AGM CD45^low^c-Kit^high^ cells led to the consistent formation of cell clusters in vitro during at least 8 passages of cocultures with stromal cells. In contrast, knock-down of Sox17 in the Sox17-transduced cells led to dispersed hematopoietic colonies containing granulocytes and macrophages in the co-culture with stromal cells [10]. These reports suggest that Sox17 contributes to the maintenance of cells with stem cell phenotypes in the hematopoietic cell clusters of the AGM region and that downregulation of Sox17 induces hematopoietic differentiation [10].

The Notch signaling pathway plays a significant role in fetal hematopoiesis. Notch1- knockout mouse embryos were found to be defective for definitive hematopoiesis and abnormal vascular formation [22]. Studies show that Sox17 interacts directly with the promoter region of Notch 1 and induces the expression of Notch1, followed by the Hes1 induction [17, 23]. The Sox17-Notch1-Hes1 axis is suggested to play a role in the maintenance of the undifferentiated state in Sox17-transduces cells [17, 23]. Moreover, the thrombopoietin (TPO) signaling pathway is important for the formation of Sox17-transduced IAHCs that highly express the TPO receptor c-Mpl and the maintenance of hematopoietic activity. The expression of c-Mpl was observed in IAHCs of E10.5 embryos and the stimulation of the TPO leads to the emergence of hematopoietic cells from the endothelial cells of the AGM region [24].

Up until now, we have shown that introduction of Sox17 in IAHC cells in the AGM region results in the formation of cell clusters with hematopoietic potential [10, 17, 23, 24, 25] and assume that Sox17 induces the expression of target genes to maintain an undifferentiated state via the interaction of Sox17 with its promoter. Such genes include Notch1 [17], ESAM, and VE-Cad [23] and each gene induced by Sox17 may have distinct functions in the maintenance of undifferentiated states. However, it remains to be determined how these genes function to confer cluster formation and hematopoietic activity by the Sox17 and whether there still are unidentified Sox17 target genes. In the present study, RNA-seq analysis was done to identify genes that are upregulated in the *Sox17*-expressing IAHCs from the 10.5 AGM region as compared with *Sox17*-nonexpressing ones, according to the method by Kayamori et al. [26] with the use of mice carrying the enhanced GFP gene in frame with Sox17 start codon [27, 28]. In this study, we focus on the *Rasip1* gene because among the top 7 highly expressed genes, Rasip1 was reported to be a vascular-specific regulator but not recognized as a hematopoietic regulator. We here further show the functional role of Rasip1 in the maintenance of HSC-containing IAHC cells of the E10.5 AGM region.

## Methods Animals

ICR mice were purchased from Japan SLC. Animal experiments were performed by institutional guidelines approved by the Animal care Committee of Tokyo Medical and Dental University (approval number: A2018-265C, A2019-108C, A2021-177C).

### Isolation of IAHCs from the AGM region

AGM regions excised from E10.5 ICR mice were incubated in 1 mg/ml Dispase II (Roche, Basel, Switzerland) for 20 min at 37 °C. After washing cells with stop solution [Hank’s balanced salt solution (HBSS) containing 10% (v/v) fetal calf serum (FCS) and 250 μg/ml DNase I (Roche)], cells were resuspended in Dulbecco’s modified Eagle’s medium (DMEM) containing 2% (v/v) FCS. The cells were subjected to immunostaining with phycoerythrin (PE)-conjugated anti-mouse CD45 (30-F11) and allophycocyanin (APC)-conjugated anti-mouse c-Kit (2B8) antibodies (eBioscience, San Diego, CA, USA). Stained cells were suspended in DMEM supplemented with 2% (v/v) FCS and 1 μg/ml propidium iodide (PI) (Calbiochem, Darmstadt, Germany) and analyzed by FACS Aria II (Becton Dickinson, Lincoln, NJ, USA). The results of flow cytometry were analyzed with FlowJo (Three Star Inc., Ashland, VT, USA). Sorted CD45^low^c-Kit^high^ cells seeded on OP9 stromal cells in an alpha-minimal essential medium (α-MEM) supplemented with 10% (v/v) FCS, 25 ng/ml stem cell factor (SCF, PeproTech, Rocky Hill, NJ), 10 ng/ml interleukin-3 (IL-3, PeproTech), and 10 ng/ml thrombopoietin (TPO, PeproTech).

### RNA-seq analysis

RNA-seq analysis was done as described previously [26]. We used the AGM region of heterozygous embryos, in which the Sox17 gene was replaced with the enhanced GFP gene [27, 28]. Total RNA was isolated from the E10.5 Sox17 (GFP)^+^ mouse AGM region-derived CD45^low^c-Kit^high^ cells treated with DMSO, 30 nM of PTC299, 200 nM of decitabine, and a combination of both agents using the RNeasy Mini Kit (Qiagen). After reverse transcription, the libraries for RNA-seq were generated from fragmented DNA using a Next Ultra DNA Library Prep Kit (New England BioLabs). After the quantification of the libraries by TapeStation (Agilent), samples were subjected to sequencing with Hiseq1500 (Illumina). RNA-seq raw reads (fastq files) were mapped to a mouse genome. Gene level counts for fragments mapping uniquely to the mouse genome were extracted from BAM files. Gene expression values were then calculated as reads per kilobase of exon units per million mapped reads using cufflinks (version 2.2.1). Table S1 shows genes expressed at high levels in Sox17 (GFP)^+^ CD45^low^c-Kit^high^ cells of E10.5 Sox17-GFP embryo as compared with Sox17 (GFP)^−^ CD45^low^c-Kit^high^ cells.

### Retroviral transduction of genes into cells

Plat-E cells for packaging ectopic retroviruses [29] were seeded 1 day before transfection. pMY vector inserted with a gene of interest was transfected into PlatE packaging cells with Trans IT-293 Reagent (Mirus, Madison, WI, USA). After 2 days of culture, the culture medium containing retroviruses was collected. CD45^low^c-Kit^high^ cells sorted from the E10.5 AGM region were infected with retroviruses encoding the Sox17-, Sox17-ERT-, Rasip1- internal ribosomal entry sequence (IRES)-GFP or the Sox17-IRES-mCherry gene in the presence of 10 μg/ml polybrene at 37 °C for 2.5 h. Transduced cells were co-cultured with OP9 stromal cells in α-MEM containing 10% FCS, 25 ng/ml SCF, 10 ng/ml IL-3 and 10 ng/ml TPO.

### Tamoxifen-dependent nuclear translocation of Sox17 and analysis of Rasip1 expression

The Sox17-ERT-IRES-GFP-encoding retrovirus was infected with CD45^low^c-Kit^high^ cells from the E10.5 AGM region [30]. The transduced cells were co-cultured with OP9 stromal cells supplemented with SCF, IL-3, and TPO with or without tamoxifen (1 μg/ml tamoxifen citrate). After 3 days of culture, Sox17-ERT-IRES-GFP-transduced cells were sorted by flow cytometry. cDNA was synthesized from total RNA isolated from sorted cells and Rasip1 expression was analyzed by RT-PCR.

### Whole-mount immunostaining

Whole-mount immunostaining was performed according to a previously described protocol [5, 17, 23]. E10.5 ICR mouse embryos were fixed in 2% paraformaldehyde (PFA) for 20 min on ice. After washing with PBS, the embryos were treated with PBS-MT [1% (w/v) skim milk powder, and 0.4% (v/v) Triton X-100] containing 2% (w/v) bovine serum albumin for 1 h on ice and then immunostained with a rat anti-mouse CD117 (c-Kit) antibody (2B8; BD Pharmingen), a goat anti-mouse CD31 antibody (Bio-Techne, Minneapolis, MN, USA), a rabbit anti-mouse Rasip1 antibody (Proteintech, Rosemont, IL, USA) overnight at 4 °C. Immunostained embryos were washed three times with PBS-MT for 1 h each and then stained with Alexa Fluor 647-conjugated donkey anti-rat IgG (Jackson Immuno Research, West Grove, PA), Alexa Fluor 546-conjugated donkey anti-goat IgG (Molecular Probes, Oregon, USA), Alexa Fluor 488-conjugated donkey anti-rabbit IgG (Life Technologies, California, USA) in PBS-MT overnight at 4 °C. After washing three times with PBS-MT for 1 h each, embryos were stained with Hoechst 33258 (Nacalai Tesque, Kyoto, Japan) in PBS-T (0.4% [v/v] Triton X-100 in PBS) for 20 min at 4 °C, followed by washing twice with PBS-T for 20 min each at 4 °C. Moreover, embryos were treated with 50% (v/v) methanol/PBS for 10 min at 4 °C and then treated twice with 100% methanol for 10 min each at 4 °C. The embryos were washed in a 50% (v/v) methanol/a mixture of benzyl alcohol and benzyl benzoate (1:2, BABB) three times. Finally, the embryos were mounted in BABB. Observations were performed by confocal laser-scanning microscopy (LSM 510; Carl Zeiss, Oberkochen, Germany).

### Luciferase assay

NIH3T3 cells (1.0 × 10^6^) were transduced with a pEFBOSE vector encoding Sox17, Sox17G103R mutant, which disrupt the DNA binding capacity, pGL3 vector (Promega, Madison, WI) carrying the putative Rasip1 promoter or point mutated Rasip1 promoters and pRL-CMV or pRL-TK encoding sea pansy luciferase genes with Trans-IT LT1 reagent (Mirus, Madison, WI). After 3 days, cells were solubilized and luciferase activities in cell lysates were monitored by the Dual-Luciferase Reporter Assay System (Promega, Madison, WI), and a Mithras LB 940 (Berthold Technologies, Bad Wildbad, Germany) was used for quantitation. First, the pRL-CMV vector (Promega, Fitchburg, WI) and Sox17G103R mutant was used in the control luciferase assay. Then, the pRL-TK vector (Promega, Fitchburg, WI) was mainly used in the luciferase assay.

### Knockdown of Rasip1 in CD45^low^c-Kit^high^ cells prepared from the E10.5 AGM region

The shRNA sequences and hairpin oligonucleotides designed with 9-mer nucleotide spacer TTCAAGAGA [31] were annealed downstream of the U6 promoter in the pMKO. shRNA sequences were as follows: 5’-ACGTGTGCTGTCCGT-3′ (shRasip1); 5’- CAAACGCTGAGTACTTCGA-3′ (shLuc). In the same retrovirus vector containing U6 promoter-shRasip1 or shLuc, a GFP gene was expressed under the control of the SV40 promoter [9]. The shRNA and GFP constructs were introduced into Sox17-IRES-mCherry transduced cells derived from the E10.5 CD45^low^c-Kit^high^ cells by a retrovirus, followed by co-culture with OP9 cells for 4 days. After 4 days of co-culture, Sox17-IRES-mCherry transduced cells formed clusters that expressed bright GFP (shRNA). And the images of these clusters were taken with fluorescent microscopy. The volume of each Mock- and shRNA-transduced cell cluster was evaluated by the formula described by Wu et al. [32]. Cluster volume = length × width^2^/2, where length represents the largest diameter and width represents the perpendicular diameter to the largest diameter. We then sorted GFP^+^ cells and assessed the colony-forming ability (2.5 × 10^3^ GFP^+^ cells) after 7 days of the culture in a semi-solid medium (Methocult M3434, Stem Cell Technologies, Vancouver, Canada) and the expression level of Rasip1 (2.0 × 10^4^ GFP^+^ cells) using RT-PCR. Rasip1 KD efficiency was assessed by Image J and expressed in percentage as compared with β-actin expression.

### Reverse transcription-polymerase chain reaction (RT-PCR)

Sorted cells were dissolved in ISOGEN (WAKO, Osaka, Japan) and RNAs were recovered according to the manufacturer’s protocol. Then, cDNA was synthesized from total RNA isolated from sorted cells. Expression values for the Rasip1-knockdown or overexpression of the *Rasip1* gene were normalized to the levels of the β-actin gene. PCR amplification was performed using rTaq (TAKARA Bio Inc., Otsu, Japan) under the following conditions: 96 °C for 3 min, and then cycles of 96 °C for 30 s, 55 °C for 30 s, and 72 °C for 30 s. The primer sets used were as follows:

5′ GTGACAGATGACGCATTGCATAGAGAA-3′;

3′-ACTCAAACTCTCAGCCGTCTACGCAGG-5′ (Rasip1),

5′-CCAGGGTGTGATGGTGGGAA-3′; 3′-CAGCCTGGCTGGCTACGTACA-5′ (b-actin).

### Overexpression of Rasip1 in CD45^low^c-Kit^high^ cells prepared from the E10.5 AGM region

A full-length cDNA encoding Rasip1 was inserted into the pMY-IRES-GFP vector with a Flag-tag. FACS-sorted CD45^low^c-Kit^high^ cells prepared from E10.5 AGM regions were infected with the retroviruses encoding the Rasip1-IRES-GFP gene in the presence of 10 mg/ml polybrene for 2.5 hrs. After 4 days of culture on OP9 stromal cells, GFP-encoding virus-infected cells were sorted by FACS Aria II and reseeded onto new OP9 stromal cells to proliferate the Rasip1-transduced cells. The process was repeated every 3 or 4 days.

### Statistical analysis

All data of luciferase activities and colony counts are represented as the mean± standard deviation. Comparisons between two samples were performed using Student’s t-tests.

## Results

### Rasip1 is expressed in endothelial cells and cell membrane site of IAHCs of the E10.5 AGM region

We have previously reported that the CD45^low^c-Kit^high^ cell population in the E10.5 AGM region has the highest hematopoietic activity among all the populations classified by CD45 and c-Kit expression levels [23]. Furthermore, we have demonstrated the forced expression of the transcription factor Sox17, which was previously suggested to maintain the hematopoietic activity in fetal hematopoiesis in E10.5 AGM CD45^low^c-Kit^high^ cells, leads to the consistent formation of cell clusters with hematopoietic activity in vitro [10]. In the present study, we first identified genes with high expression levels in Sox17 (GFP)^+^ CD45^low^c-Kit^high^ cells as compared with Sox17 (GFP)^−^ CD45^low^c-Kit^high^ cells of the AGM regions in E10.5 Sox17 promoter-GFP knock-in embryos using RNA-seq analysis. Table 1 indicates the top 7 highly expressed genes that are likely to be transcriptional targets of Sox17. Within these top 7 genes that are highly expressed in Sox17 (GFP)^+^ CD45^low^c-Kit^high^ cells, the *Rasip1* is the least studied gene in murine embryonic hematopoiesis. Therefore, and because Rasip1 was reported to be a vascular-specific regulator, we focused on the involvement of the *Rasip1* gene in the formation of cluster cells in the E10.5 AGM region. Rasip1 is a protein that acts as a vascular-specific regulator of GTPase signaling, cell architecture, adhesion, and tubulogenesis [33-36]. As shown in Fig. 1A (lower panel), Rasip1 was expressed in endothelial cells and CD45^low^c-Kit^high^ cells.

**Table 1:** Top 7 selected genes expressed at high levels in Sox17 (GFP)^+^ CD45^low^c-Kit^high^ cells of E10.5 Sox17-GFP embryo as compared with Sox17 (GFP)^−^ CD45^low^c-Kit^high^ cells. FPKM: fragments per kilobase of transcript per million mapped reads. In this table, “(FPKM+1)” represents genes whose expression is 1 FPKM or more. As low as FPKM+1 was verifiable by RT-PCR and variability in RNA-Seq data greatly increases the lower expression.

|  |  | Sox17<sup>+</sup>: (FPKM+1)/<br>Sox17<sup>-</sup>: (FPKM+1) |
| --- | --- | --- |
| 1 | *Cldn5* | 242.21 |
| 2 | *Sox18* | 126.71 |
| 3 | *Aplnr* | 121.16 |
| 4 | *Esam* | 87.78 |
| 5 | ***Rasip1*** | 75.09 |
| 6 | *Plvap* | 69.84 |
| 7 | *Emcn* | 62.70 |

**Fig. 1:**
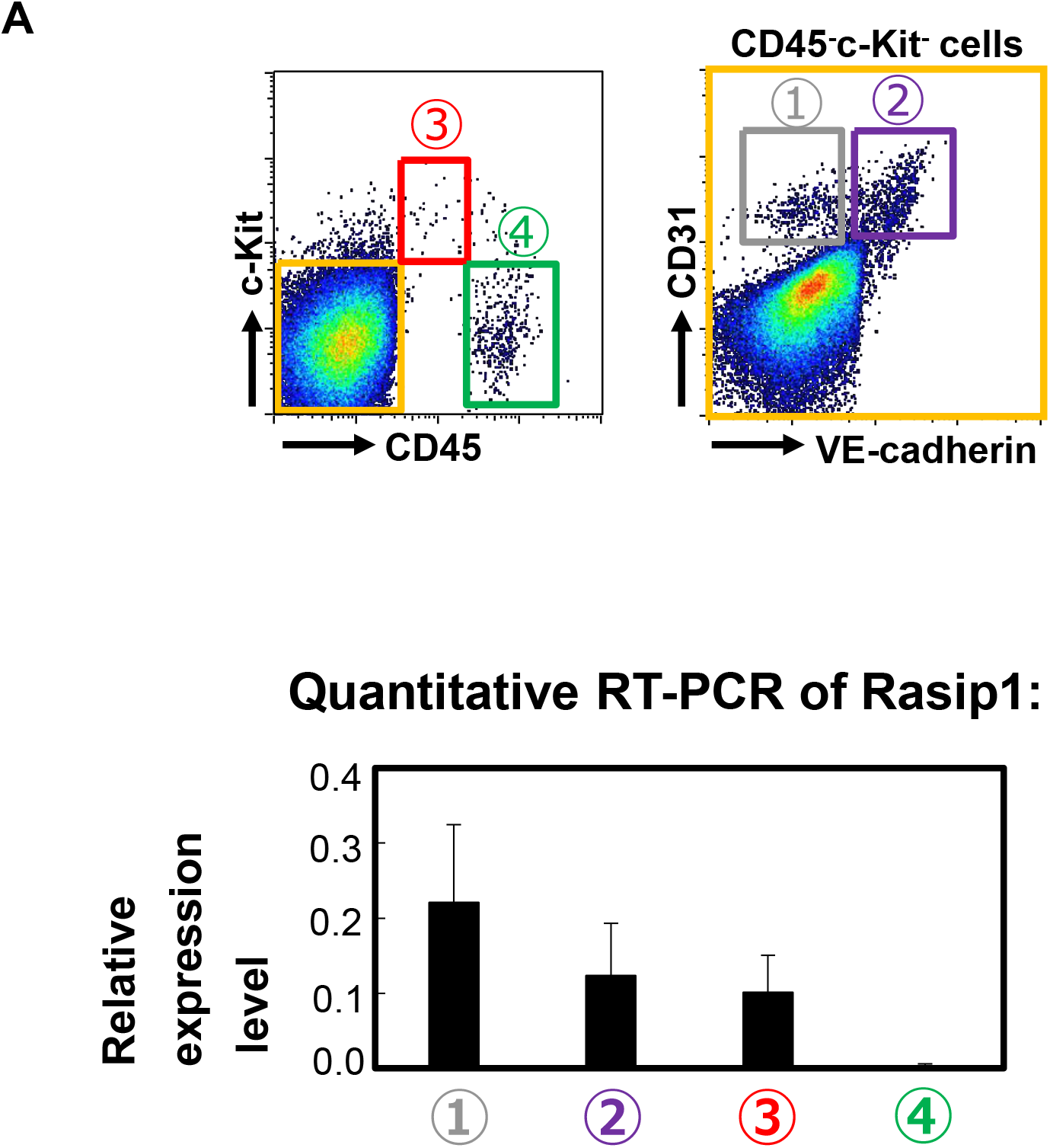
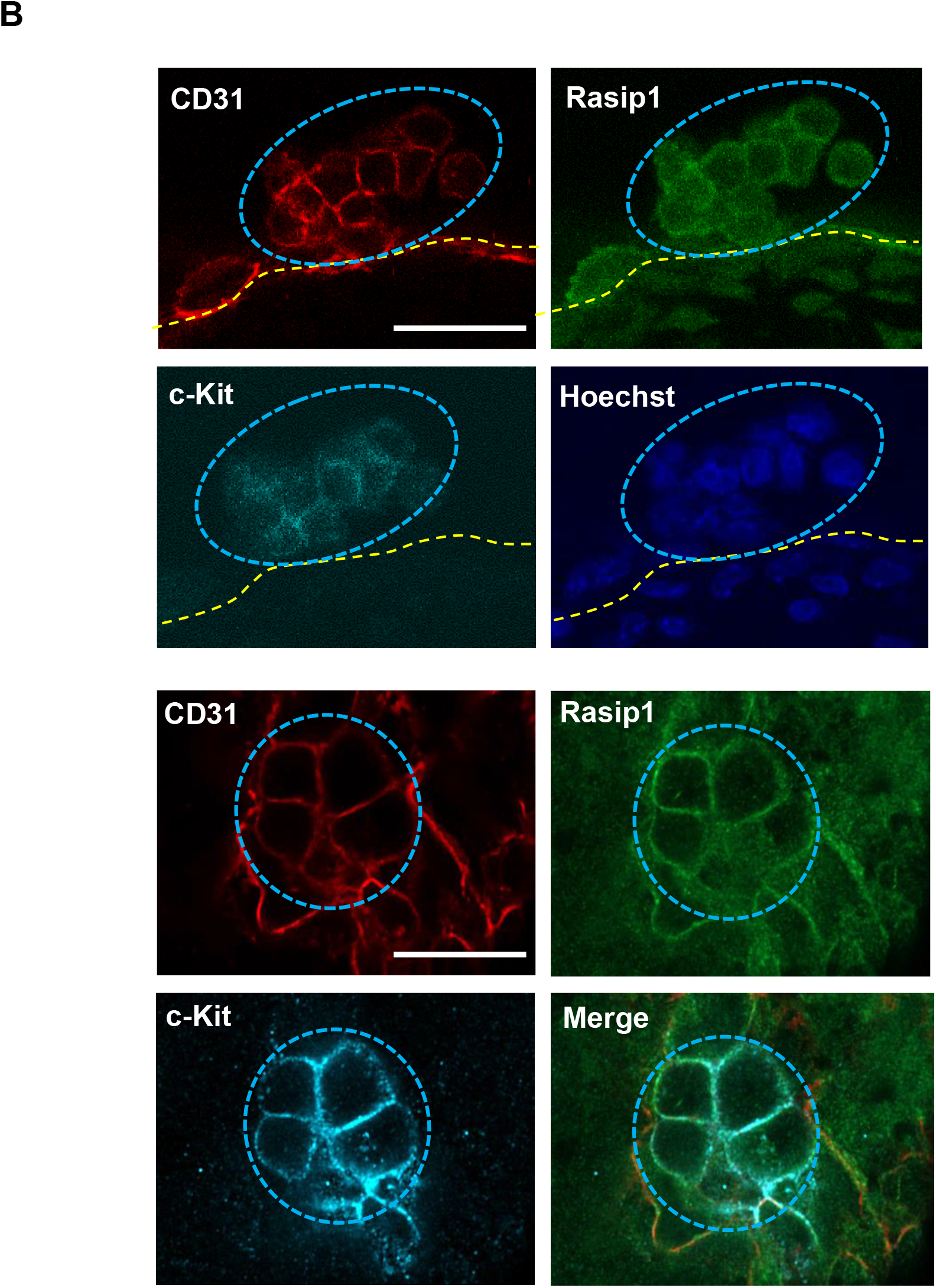
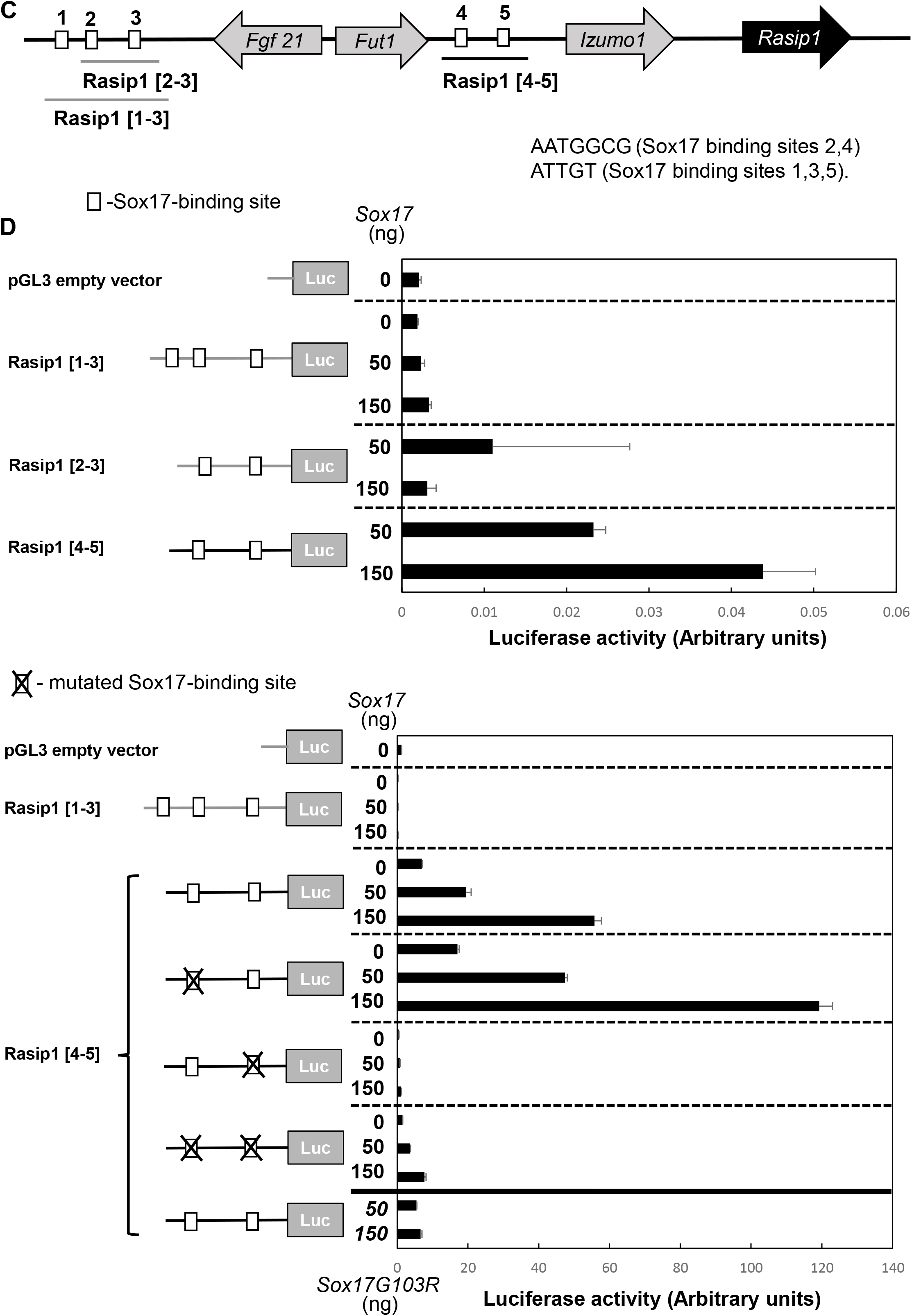
Rasip1 expression in IAHCs in the E10.5 AGM region. **A**. Upper panel: The expression profile of CD45, c-Kit, CD31, and VE-Cad in the E10.5 AGM region. The numbers from 1 to 4 indicate cell populations separated by the expression profile: 1.CD45^−^ c-Kit^−-^ CD31^+^ VE-cad^−^ cells (endothelial cells); 2.CD45^−^ c-Kit^−^ CD31^+^ VE-cad^+^ cells (endothelial cells); 3.CD45^low^c-Kit^high^ cells (containing HSPCs); 4.CD45^+^ c-Kit^−^ cells (hematopoietic cells). Lower panel: RT-PCR shows the relative expression levels of *Rasip1* in these 4 populations. **B**. Whole-mount immunofluorescence staining of Rasip1 (green), CD31 (red), and c-Kit (cyan) in IAHCs of the E10.5 AGM region. The Rasip1 expression on the membrane of IAHCs. The nuclei were stained by Hoechst 33258 (blue). Scale bar, 20 μm. **C**. A genetic scheme showing the location of the *Rasip1* gene and upstream genes in mouse chromosome 7. 5 putative Sox17-binding sites were indicated by squares with numbers 1 through 5. **D**. Upper panel: Requirement of the proximal putative Sox17-binding site (5) for induction of the Rasip1 gene by Sox17. NIH3T3 cells (1 × 10^6^) were transfected with a pGL3 vector containing the putative Rasip1 promoter sequences. pRL-CMV was co- transfected as an internal control. The region Rasip1 [1-3] (979 bp) has three putative Sox17- binding sites (1, 2, and 3), and the region Rasip1 [2-3] (602 bp) has two putative Sox17 binding sites (2 and 3). The region Rasip1 [4-5] (720 bp) has putative Sox17-binding sites 4 and 5. Two types of Sox17-binding sites are shown in the promoter regions: 1. AATGGCG (Sox17 binding sites 2, 4); 2. ATTGT (Sox17 binding sites 1,3,5). Lower panel: Inhibition of the Sox17-induced *Rasip1* gene activation by a mutation in the putative Sox17-binding site 5. NIH3T3 cells (1 × 10^6^) were transfected with a pGL3 vector containing the putative Rasip1 promoter sequences with point mutations in the putative Sox17-binding sites 4 or 5, or both together with plasmids encoding wild-type (WT) Sox17 or Sox17G103R which lost the DNA binding ability. In this experiment, pRL-TK was co-transfected as an internal control. A solid black line indicates the Rasip1 [4-5] region, solid grey lines show the pGL3 vector and Rasip1 promoter region [1-3] as controls.

We have previously demonstrated the expression of Sox17 in basal cells of the E10.5 AGM region’s IAHCs by whole-mount in situ hybridization, and CD45^low^c-Kit^high^ cells transduced with Sox17 maintain their undifferentiated state [4]. Rasip1 expression in IAHCs of wild-type (WT) embryos was examined by immunohistochemistry. As shown in Fig. 1B, immunoreactivity for Rasip1 was detected in the membrane of c-Kit^+^ IAHC cells along with that for the endothelial cell marker CD31 expression. We observed Rasip1 expression in the aortic endothelium of the AGM region as well as the IAHC cell membrane site.

To evaluate the effect of Sox17 on the *Rasip1* activation, we first searched for the putative Sox17-binding site in the upstream region of the *Rasip1* gene. Five putative Sox17- binding sites were observed in the region (Fig. 1C). We examined the luciferase activity of the promoter constructs, in which Rasip1[1-3] contained the first three putative Sox17- binding sites, Rasip1[2-3] included the second and the third putative Sox17-binding sites, and Rasip1[4-5] had the other two putative Sox17-binding sites, with the varying amount of vector encoding the Sox17 or its mutant with no DNA binding capacity (Fig.1C). As shown in Fig. 1D (upper panel), the introduction of Sox17 led to increased Rasip1 promoter activity of the Rasip1 [4-5] construct in a dose-dependent manner. In contrast, the Sox17G103R mutant, in which Gly^103^ is replaced with Arg to disrupt its DNA binding activity, did not induce the luciferase activity of the putative *Rasip1* promoter region, Rasip1[4-5] (Fig. 1D, lower panel, the bottom pair of bars). Moreover, the Rasip1 promoter activity of the Rasip1 [4-5] construct decreased abruptly when Sox17 protein was absent or had a non-functioning mutation, or a putative Sox17 binding site 5 in the *Rasip1* gene was mutated (Fig. 1D, lower panel). These data suggest that Sox17 binds to the *Rasip1* promoter region and induces expression of the *Rasip1* gene.

### Rasip1 is expressed in Sox17-transduced CD45^low^c-Kit^high^ cells with high hematopoietic activity

We previously showed that when CD45^low^c-Kit^high^ cells sorted from E10.5 AGM regions were transduced with Sox17-IRES-GFP virus and co-cultured with OP9 stromal cells with SCF, IL3, and TPO, the Sox17-transduced cells formed cell clusters with high hematopoietic activity [4]. We thus examined the expression level of Rasip1 in Sox17-transduced CD45^low^c- Kit^high^ cells by RT-PCR analysis. As shown in Fig. 2A, the *Rasip1* gene was clearly expressed in Sox17-transduced cells, while its expression was almost negligible in the Mock-transduced cells. This result indicates that the *Rasip1* gene is a target of Sox17 and implies that Rasip1 may contribute to the Sox17-transduced cell cluster formation.

**Fig. 2:**
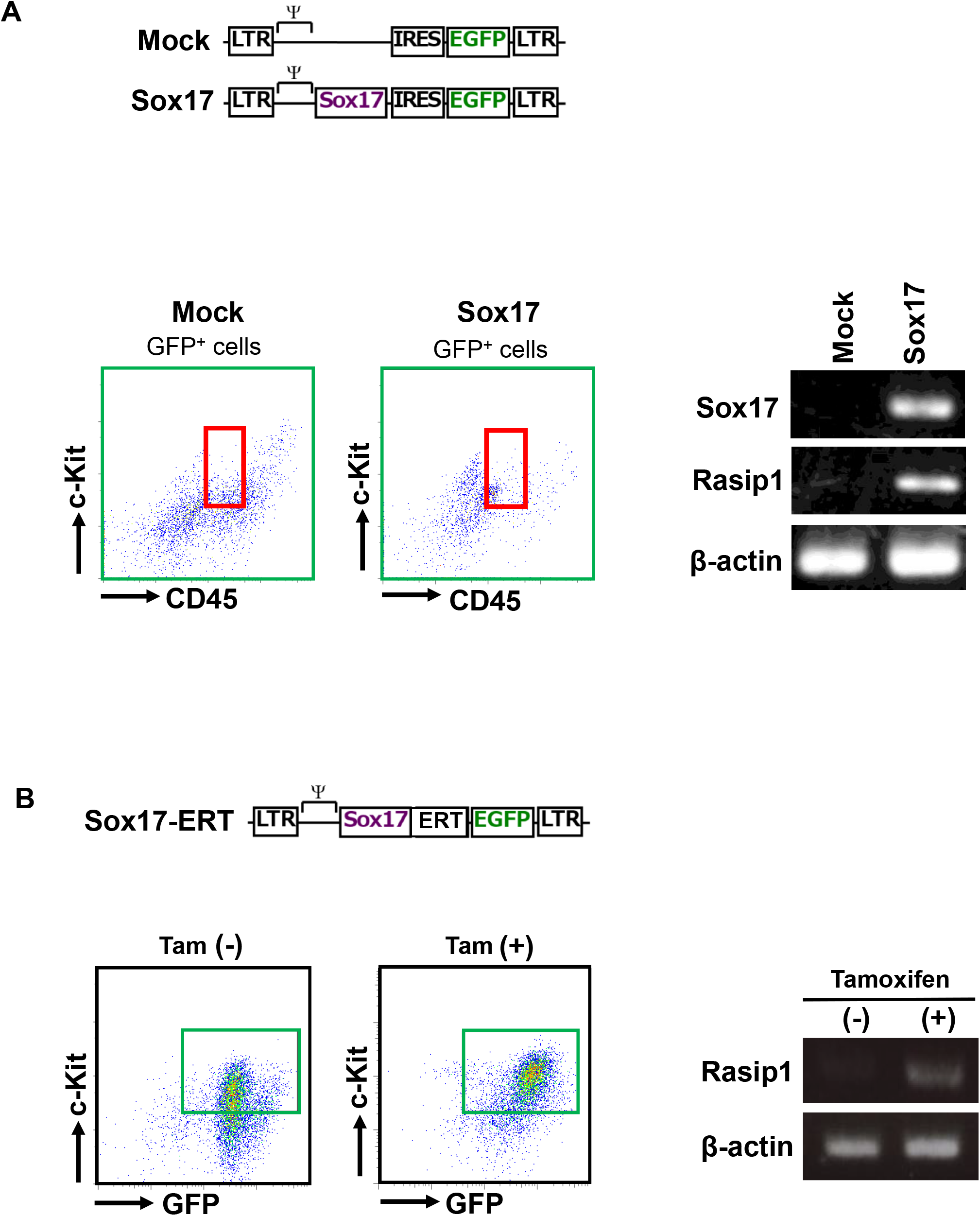
Expression of Rasip1 in Sox17 (GFP)-transduced CD45^low^ c-Kit^high^ cells. **A**. Upper panel: A schema of Sox17-IRES-GFP (Sox17) and IRES-GFP (Mock) retrovirus vectors. Lower panel: The red squares show a region of the CD45^low^c-Kit^high^ cell population. The expression of Rasip1 in Mock- or Sox17-transduced cells was analyzed by RT-PCR and normalized by β-actin. **B**. Upper panel: A schema of the Sox17-ERT-IRES-GFP retrovirus vector. Lower panel: The dark green squares show Sox17 (GFP)^+^ CD45^low^c-Kit^high^ cells. After FACS sorting, expression analysis of the *Rasip1* gene was performed in Sox17-ERT- IRES-GFP-transduced cells with or without tamoxifen (1.0 μg/ml).

Next, we examined the *Rasip1* expression level in Sox17-ERT-transduced CD45^low^c- Kit^high^ cells sorted from E10.5 AGM by RT-PCR analysis. Tamoxifen is a reagent that dose-dependently transports the Sox17-ERT fusion protein into the nucleus of the cells. As shown in Fig. 2B, tamoxifen-dependent nuclear import of Sox17-ERT proteins in the CD45^low^c- Kit^high^ cells led to an increased expression level of Rasip1 compared to control. Together, these results are in agreement with those of the luciferase assay, implying that the Sox17 transcription factor is tightly associated with the induction of Rasip1 expression.

### Rasip1 knockdown in Sox17-transduced cluster cells impedes cluster formation and decreases the hematopoietic activity

To confirm the contribution of Rasip1 to the formation of cell clusters with high hematopoietic activity, retroviruses encoding shRNA against luciferase (shLuc) and Rasip1 (shRasip1) and a GFP gene were infected with Sox17-transduced cells. After 4 days of the culture with the stromal cells, we found that most of the cells express mCherry, i.e. Sox17 (Fig. 3A, lower left panel). This is because, without Sox17, AGM cells lose the ability to maintain cluster-forming cells [10]. Among the mCherry^+^ (Sox17^+^) cells, we observed to some extent GFP^+^ cells (shLuc- or shRasip-transduced cells) (Fig. 3A, lower right panel). Required numbers of GFP^+^ cells were sorted by FACS as presented in Fig. 3B, for the analysis of Rasip1 expression in shLuc- and shRasip1-transduced cells and/or for the colony-forming assays, respectively. The expression level of *Rasip1* was decreased by 42% in cells transduced with shRasip1 as assessed by Image J (Fig.3B, right panel). Moreover, we noted that Sox17-transduced cells with downregulated Rasip1 expression by the introduction of shRNA displayed a decrease in their total and multilineage colony-forming activities (Fig. 3C). These results indicate that cells with knockdown of Rasip1 decreased the hematopoietic capacity in vitro.

**Fig. 3:**
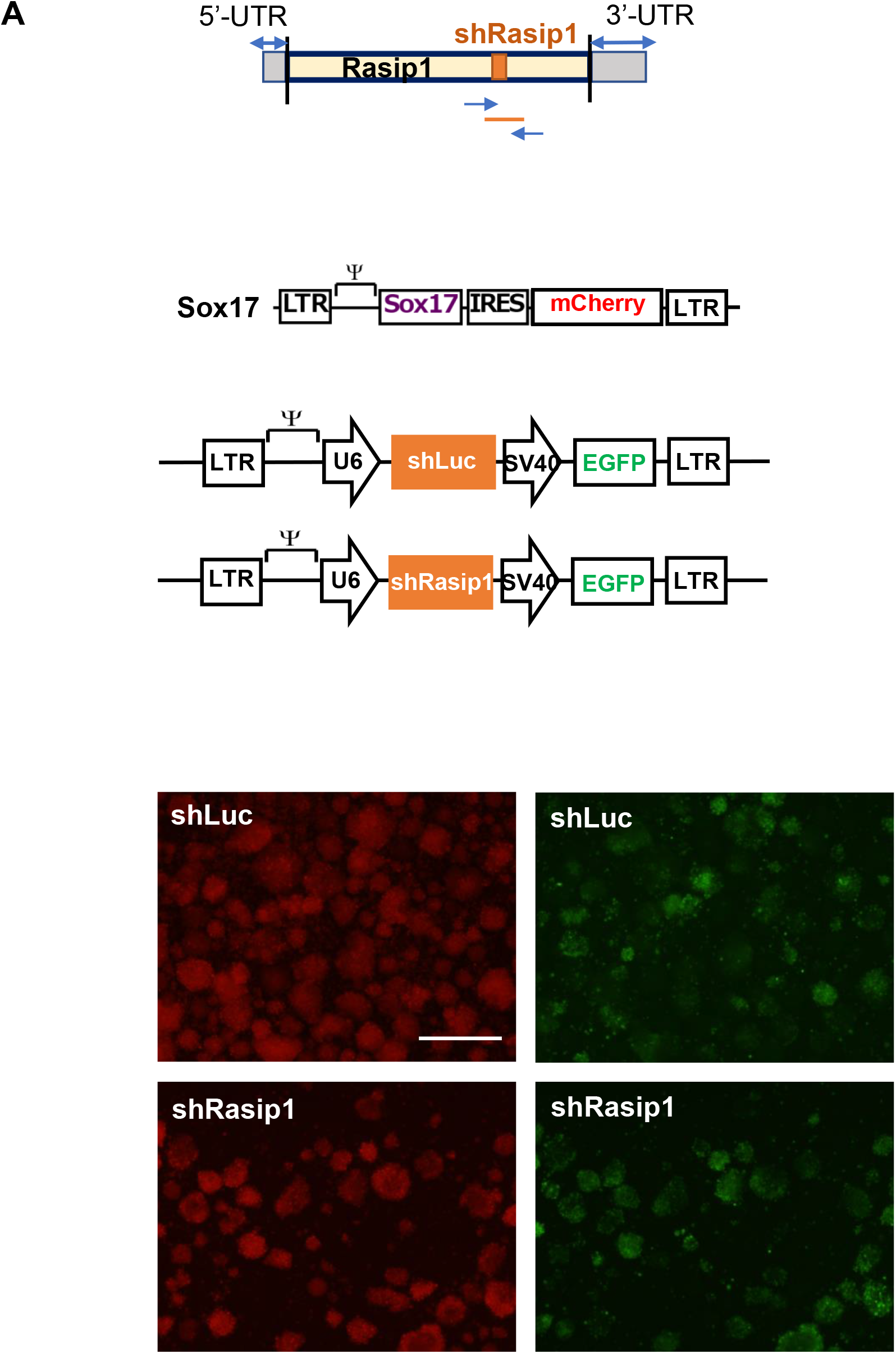
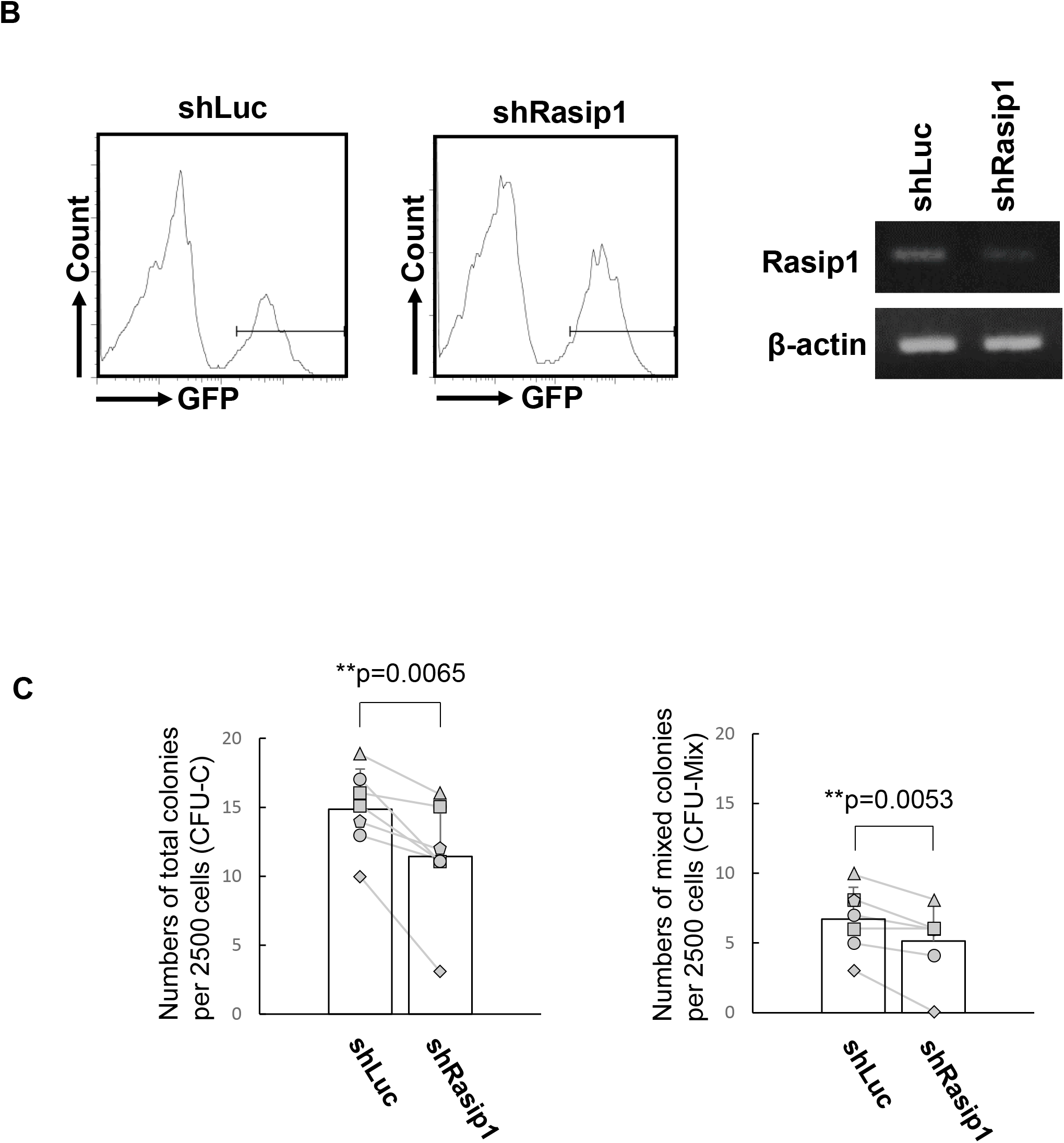
Rasip1 knockdown (KD) in CD45^low^c-Kit^high^ cells. **A**. Upper panel: A schema of the position of the Rasip1 shRNA in the Rasip1 mRNA, a retrovirus vector of Sox17-IRES- mCherry, and retrovirus vectors of shLuc and shRasip1. Lower panel: After 4 days of Sox17- IRES-mCherry transduction, Sox17-transduced cells formed cell clusters. shLuc and shRasip1 were introduced into Sox17-transduced cells and the cells were co-cultured with OP9 stromal cells. Morphologies of shRasip1 (green, GFP^+^)-transduced Sox17^+^ cells (red, mCherry^+^). Scale bar, 1.0 mm. **B**. shLuc- or shRasip1-transduced GFP^+^ cells were recovered by FACS. Expression of Rasip1 in shLuc- and shRasip1-transduced cells was analyzed by RT-PCR and normalized by β-actin gene expression. RT-PCR result is in the static image that is generated by a mirror-reversal of an original across a horizontal axis, due to the sample order in the electrophoresis. **C**. Sorted GFP^+^ cells (2.5 × 10^3^) were embedded in a semi-solid medium. The number of total (CFU-C) and mix (CFU-Mix) colonies were scored after 7 days of culture. The pair of colony numbers (sh-Luc and shRasip1) in each experimental group were plotted and connected by a line. The different marks indicate the different individual embryos (n = 7, 7 individual liters of mice from which each experimental group was set up).*p≤0.05; **p≤0.01; ***p≤0.005.

We noticed that Rasip1-GFP expressing cells tended to form smaller clusters compared with control (Fig. 4A, left two panels). Hence, we calculated and compared the average volume of these clusters (Fig. 4A, right panel). Interestingly, the mean volume of shRasip1-transduced clusters was significantly lower than that of shLuc-transduced clusters. This observation supports the notion that Rasip1 is involved in the formation of cell clusters. Next, we investigated whether the GFP intensity is correlated with Rasip1 knockdown efficiency. We sorted shLuc-GFP and shRasip1-GFP cells with relatively high GFP expression levels as shown in Fig. 4B (left two panels). The sorted cells were subjected to RT-PCR analysis of Rasip1 transcripts. As shown in Fig. 4B (right panel), Rasip1 expression was decreased by 36% in the sorted shRasip1-GFP high expression cells by Image J. In contrast, as shown in Supplementary Fig. 1 (right panel), the sorted shRasip1-GFP low expression cells showed only an 8.8% decrease in the Rasip1 expression at the transcription level as assessed by Image J.

**Fig. 4.**
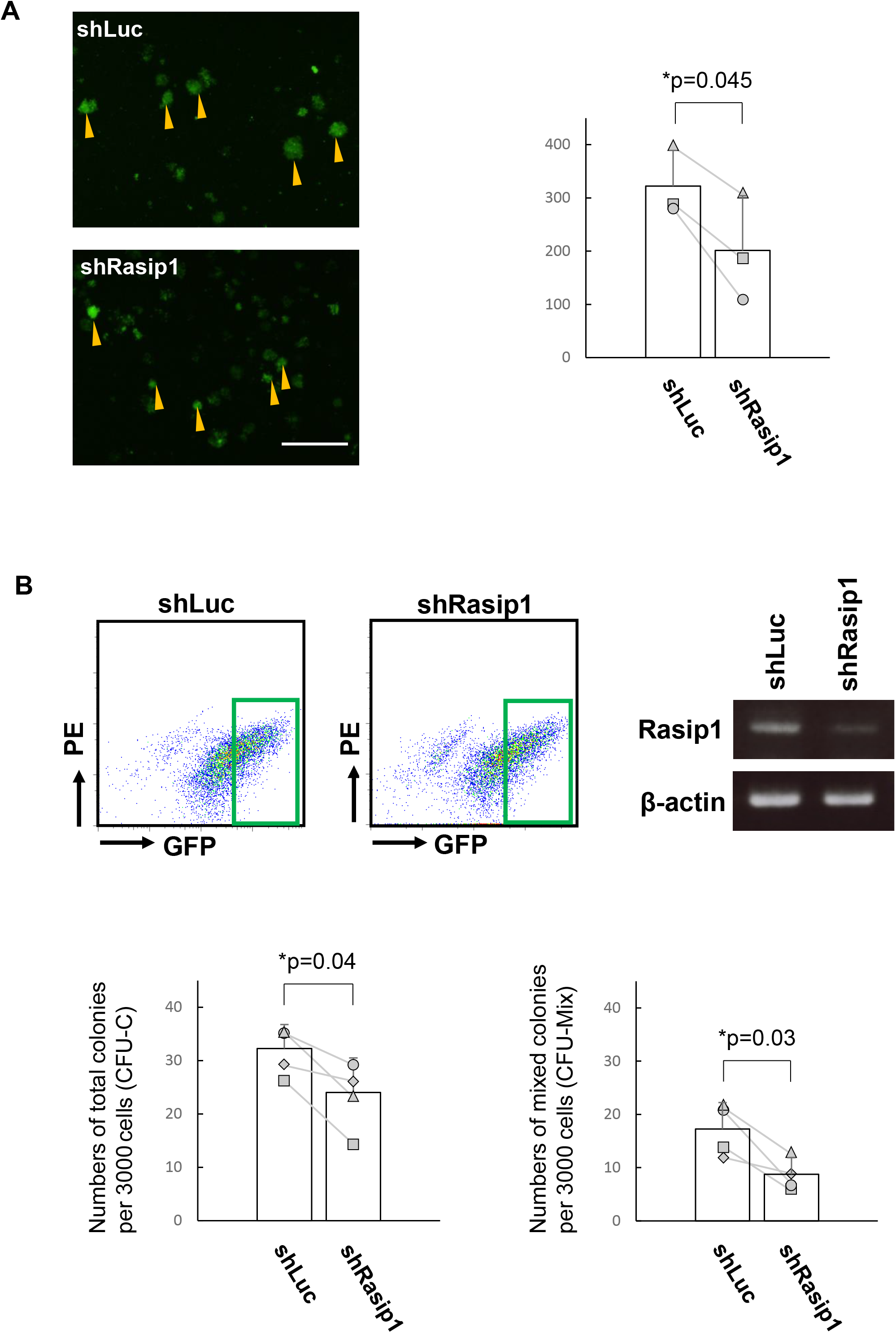
A Morphologies of GFP^+^ cell clusters. Yellow arrowheads indicate cell clusters which expressed in bright green color (GFP^+^ with high intensity). The bar graph shows the average volume and standard deviation (n=3, 3 individual liters of mice from which each experimental group was set up). Scale bar, 200 μm. The average volume of GFP^+^ cell clusters with high intensity in each group (shLuc and shRasip1) was measured and analyzed as follows: 1) we measured the longest diameter and perpendicular diameter with the longest diameter 2) we calculated the average volume as follows: Cluster volume= length x width^2^/2, where length represents the largest cluster diameter and width represents the perpendicular diameter with the longest diameter 3) we calculated the average volumes of 10 clusters in maximum in one individual experiment; 4) we calculated the total average volume of all 3 experiments (n=3, 3 individual liters of mice from which each experimental group was set up). **B**. shLuc- or shRasip1-transduced GFP^+^ cells were recovered by FACS. As indicated in the FACS profile, the dark green squares show the GFP^+^ cell population with high intensity. Expression of Rasip1 in shLuc- and shRasip1-transduced cells was analyzed by RT-PCR and normalized by β-actin gene expression. RT-PCR result is in the static image that is generated by a mirror-reversal of an original across a horizontal axis, due to the sample order in the electrophoresis. **C**. Sorted GFP^+^ cells (3.0 × 10^3^) were embedded in a semi-solid medium. The number of total (CFU-C) and multilineage (CFU-Mix) colonies were scored after 7 days of culture. The pair of colony numbers (sh-Luc and shRasip1) in each experimental group were plotted and connected by a line. The different marks indicate the different individual embryos (n=4, 4 individual liters of mice from which each experimental group was set up). *p≤0.05; **p≤0.01.

### Rasip1 overexpression in CD45^low^c-Kit^high^ cells increases the hematopoietic activity

To assess the involvement of Rasip1 in the hematopoietic activity of IAHCs (Fig. 5A), we tested the hematopoietic activity of Rasip1 overexpressing cells in vitro. We transduced CD45^low^c-Kit^high^ cells with retrovirus encoding the IRES-GFP gene (Mock) or the Rasip1- IRES-GFP gene (Rasip1) and co-cultured them with OP9 stromal cells (Fig. 5B, Day 0). After 4 days of co-culture, we observed on the panel 2 populations, (1) CD45^low^c-Kit^high^ and (2) CD45^low^c-Kit^low^ populations, the former of which was missing in mock-transduced cells (Fig. 5C, left two panels). Cells indicated in red squares in Fig. 5C (left two panels) were embedded in the semisolid medium for colony-forming assays. As shown in Fig. 5D top two panels (Colony assay 1), Rasip1-transduced cells displayed a significantly higher ability to form total colonies (CFU-C) and mixed colonies (CFU-Mix) compared with Mock-transduced cells. Next, we examined whether this ability was maintained during the subculture of Rasip1-transduced cells with OP9 cells. In the subculture, after 8 days from the initial culture, cells indicated in red squares in Fig.5C (right two panels) were sorted and subjected to colony-forming assays. As shown in Fig. 5D lower two panels (Colony assay 2), the number of total colonies (CFU-C) of Rasip1-transduced cells was significantly larger than that of the mock control, while the number of mixed colonies (CFU-Mix) of Rasip1-transduced cells was statistically insignificant as compared with the mock control. Cells transduced with Rasip1 maintained their multipotent colony-forming activity for at least 7 days. Therefore, these results indicate that *Rasip1*, a Sox17 downstream target gene, endows Sox17-untransduced CD45^low^c-Kit^high^ AGM cells with the capacity to maintain hematopoietic activity, although it is transient.

**Fig. 5:**
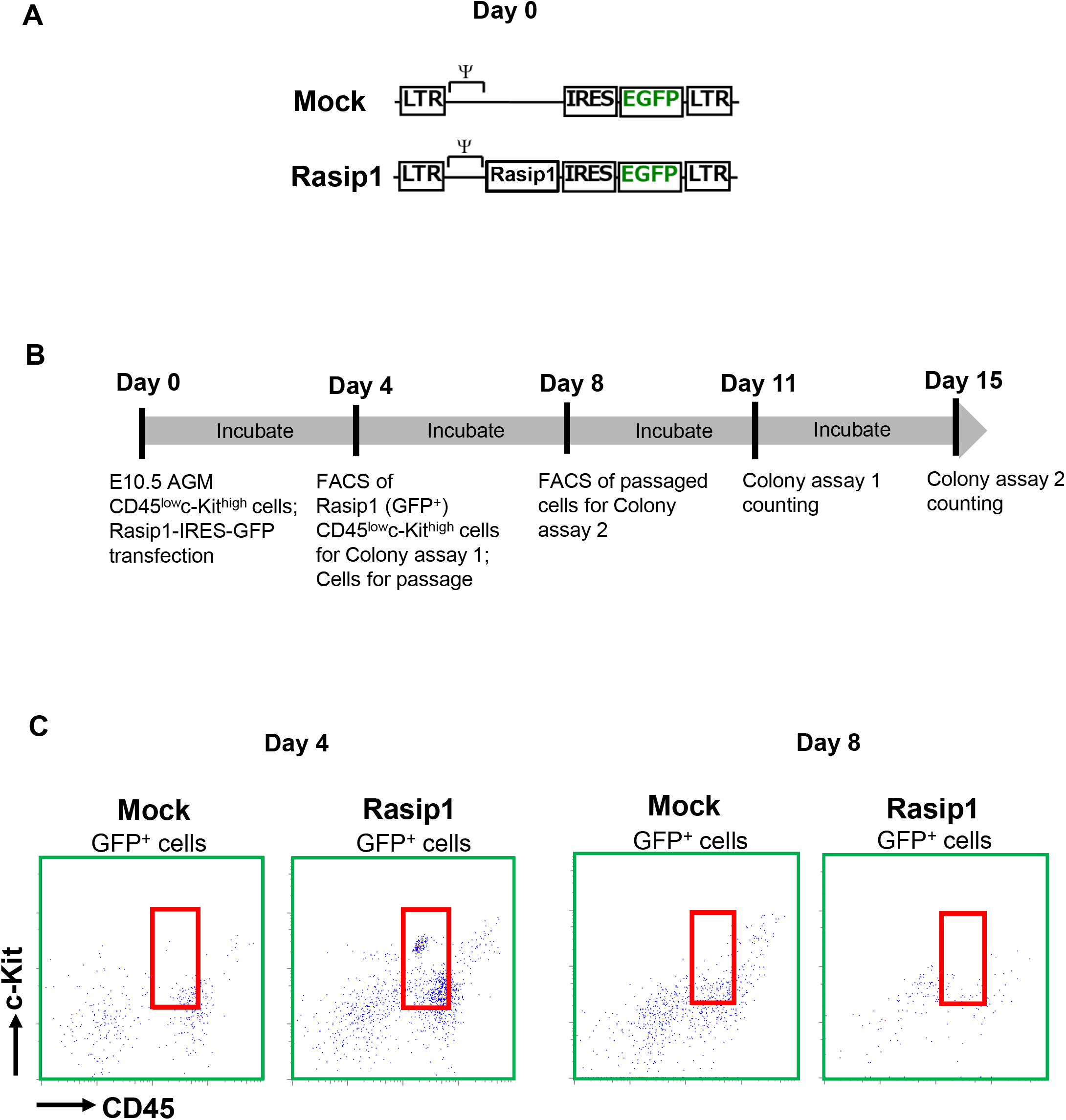
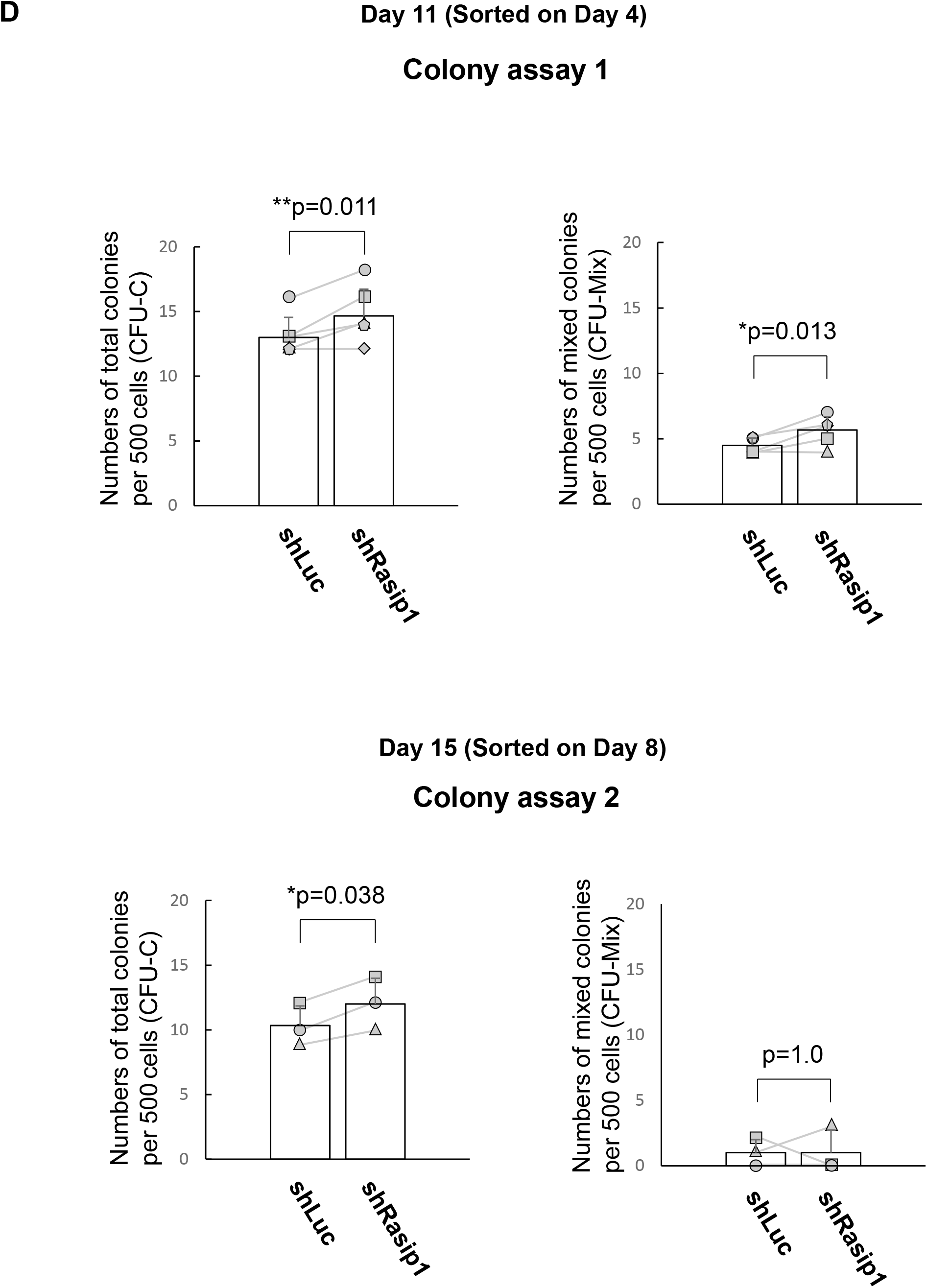
Rasip1 overexpression (OE) in CD45^low^c-Kit^high^ cells. **A**. A schema of IRES-GFP (Mock) and Rasip1-IRES-GFP retrovirus vectors. **B**. Timeline of Rasip1 OE experiment. **C**. Red squares show Mock- and Rasip1-transduced CD45^low^c-Kit^high^ cells. **D**. After 4 days of Mock and Rasip1 transduction, CD45^low^c-Kit^high^ cells (2.5 × 10^2^) were embedded in a semi- solid medium (Colony assay 1). According to the timeline in Fig.5B, CD45^low^c-Kit^high^ cells were sorted (5.0 × 10^2^) after 8 days of Rasip1 transduction (Colony assay 2). The number of total (CFU-C) and multilineage (CFU-Mix) colonies were scored after 7 days of culture (n=5). The pair of colony numbers (Mock and Rasip1) in each experimental group were plotted and connected by a line. The different marks indicate the different individual embryos (n=6, 6 individual liters of mice from which each experimental group was set up). *p≤0.05; **p≤0.01.

## Discussion

This report provides the first evidence that the *Rasip1* is a novel Sox17 downstream target gene and proposes the idea that Rasip1 contributes to the maintenance of the property of IAHCs of the E10.5 AGM region. We showed that Rasip1 is expressed in the IAHCs along with endothelial cells in immunostaining. This may be related to a previously reported function of Rasip1, a vascular-specific regulator [34-36], which is interesting in the sense that HSCs arise from the hemogenic endothelium of the dorsal aorta [10]. Rasip1 is expressed in Sox17-transduced CD45^low^c-Kit^high^ cells which contain HSC and HPC. From the luciferase assays, Sox17-induced Rasip1 expression requires the most proximal Sox17 binding site of the *Rasip1* gene promoter. Functionally, downregulation of Rasip1 in Sox17-transduced CD45^low^c-Kit^high^ cells leads to the inhibition of the multilineage colony-forming ability, while upregulation of Rasip1 causes the increase of the colony-forming ability. Our study indicates the idea that Sox17 positively regulates hematopoiesis at least in part through Rasip1 expression in IAHCs, which arises from endothelial cells as depicted in Fig. 6.

**Fig. 6:**
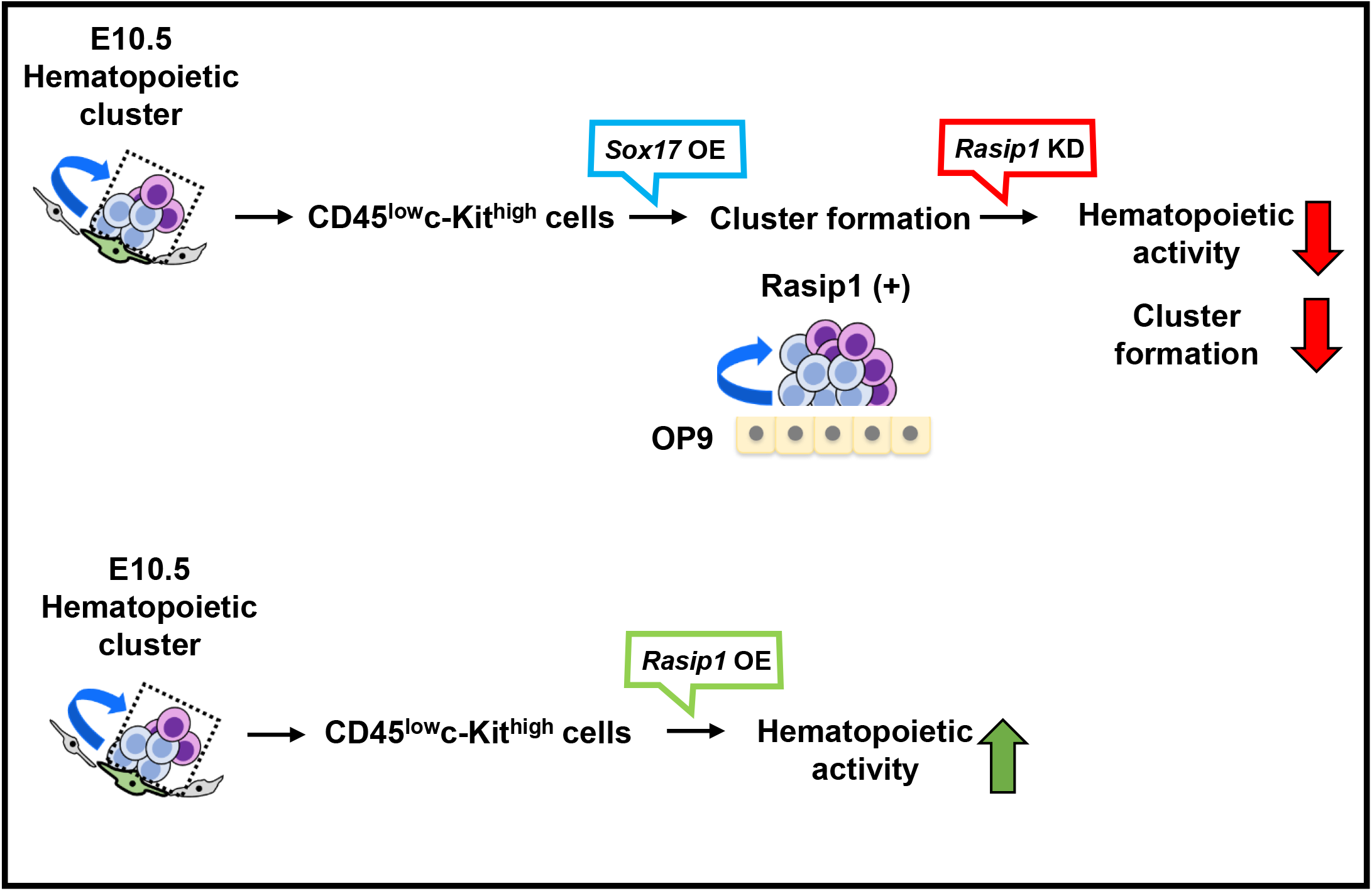
Cluster formation of IAHCs about Rasip1. A model of Rasip1 function in IAHCs. Sox17 induces the expression of Rasip1 which eventually leads to the hematopoietic activity of HSCs in IAHCs.

Rasip1 was reported as an endothelial cell-specific regulator of GTPase signaling, cell architecture, and adhesion, which is essential for endothelial cell morphogenesis and blood vessel tubulogenesis [34-38]. Mitin et al. reported the identification of Rasip1 which displays the characteristics of an endomembrane Ras effector [39]. Their experiments showed that Rasip1 possesses a Ras-associating domain (RA), which is homologous to the RA domains of other Ras “effectors”, and that Rasip1 binds to the GTP-loaded form of Ras, both in vitro and in vivo [39]. Components of the Ras signaling pathway are important during embryonic vasculogenesis and angiogenesis [39]. Xu et al. reported that the *Rasip1* gene is significantly expressed in vascular endothelial cells throughout development [38]. *Rasip1* expression is undetectable in *VEGFR2* null embryos, which lack endothelial cells, suggesting that *Rasip1* is specifically expressed in endothelial cells [38]. Rasip1 is essential for embryonic vessel formation and maintenance, but not in adult vessels [37]. Together, these data suggest that the *Rasip1* is an endothelial factor that plays a role in vascular development. Rasip1 continued to be expressed in the endothelium of vessels into the postnatal stages and was detected in adult organs, particularly in the vascularized lung [39]. Previous studies reported that Rasip1 knockout mice had formed patent lumens [36]. At E9.0, *Rasip1*^−/−^ embryos exhibited a small size, pale, hemorrhage, and pericardial edema [34]. At E10.5, *Rasip1*^*−/−*^ embryos were markedly smaller than control littermates with severe edema and hemorrhage. Viable *Rasip1*^−/−^ embryos were not detected past E10.5 [36]. Taken together, they concluded that targeted disruption of mouse *Rasip1* results in abnormal cardiovascular development and midgestational lethality [36]. From these data, we predict that the absence of Rasip1 in vivo disrupts the development of blood vessels including the dorsal aorta of the AGM region where the hemogenic endothelial cells reside, and will eventually prevent the emergence of hematopoietic clusters from hemogenic endothelium and furtherly fail the development of embryonic hematopoiesis.

The IAHC cells express adherent molecules, which are VE-Cad and CD31, on the membrane side of the endothelial cells in the dorsal aorta [5, 23]. In Fig. 1B, whole-mount immunostaining images show that IAHCs express CD31 and Rasip1 on the membrane of IAHC cells. In Table 1, other top selected genes such as endothelial cell-specific *Emcn, Plvap, Cldn5* which are involved in blood vessel development or tight junction, and *ESAM* which is involved in adhesion molecule activation are targets of Sox17 transcription factor. The Rasip1 appears to regulate endothelial cell-cell junction in a mode different from that of cell adhesion molecule, suggesting that Rasip1 alters junctional actin organization [36]. Post et al revealed that Rap1 is associated with a small GTPase regulating cell-cell adhesion, cell-matrix adhesion, and actin rearrangements, and all processes dynamically coordinate during cell spreading and endothelial barrier function [32]. At cell-cell contacts, Rasip1, in conjunction with the Rap1 effector ras-association and dilute domain-containing protein (Radil), suppresses RhoA/ROCK signaling via the RhoA, RhoGAP, and ArhGAP29 [33, 35]. The Rasip1–ArhGAP29 pathway functions in Rap1-mediated regulation of endothelial junctions, which controls endothelial barrier function [30]. In this process, Rasip1 cooperates with its close relative Radil to inhibit Rho-mediated stress fiber formation and induce junctional tightening. [33]. Moreover, it was reported that Heart of Glass 1 (HEG1), a transmembrane receptor that is essential for cardiovascular development, binds directly to Rasip1. Rasip1 localizes forming endothelial cell (EC) cell-cell junctions and silencing HEG1 prevents this localization. These studies established that the binding of HEG1 to Rasip1 mediates Rap1-dependent recruitment of Rasip1 to and stabilization of EC cell-cell junctions [35]. HEG1 is required for Rasip1 translocation to cell-cell contacts. However, the formation of a Rasip1-Rap1 or Rasip1-Radil-ARHGAP29 complex is not dependent on the presence of HEG1. From these reports, we speculate that Rasip1 has a regulatory effect on cluster integrity as a Sox17-downstream regulator of cell adhesion and cell-cell junction.

We already know that Rasip1 is endothelial-specific and essential for embryonic vascular development [36, 37]. In addition to this previous report, we found out that Rasip1 has a role in contributing to maintaining the hematopoietic activity and cluster formation of Sox17-transduced CD45^low^c-Kit^high^ cells. As shown in Fig. 4A, we have assessed the volume of GFP^+^ cluster cells with high intensity. From the result, the volume of clusters of cells with high GFP intensity in the Rasip1 knockdown group was decreased when compared with the control. In previous reports, Sox17-transduced cells highly expressed adherent molecules to form cell clusters and genes associated with the hematopoietic capacity to maintain the undifferentiated state of HSC-containing cluster cells [4]. In the scheme of mouse chromosome 7 where the *Rasip1* is located (Fig. 1C), we observed the putative Sox17 binding site (5), which is found to be located downstream before the *Izumo1* gene that encodes Izumo1 protein, activates the *Rasip1* gene expression. Functionally, Izumo1 has a different physiological role than Rasip1. In mammalian fertilization, Izumo1 binds to its egg receptor counterpart, Juno also known as folate receptor 4, to facilitate recognition and fusion of the gametes. Still, there were no reports of the interplay between the *Izumo1* gene and *Sox17* transcription factor in RNA sequence data of highly expressed genes of the Sox17. Additionally, our study shows that the *Rasip1* upregulation in vitro maintains IAHCs in an undifferentiated state, although for a short period in vitro. However, it is not fully understood how Rasip1 contributes to the hematopoietic activity of IAHC cells and what factors cooperate with Rasip1 for long-term maintenance of such activity.

In this study, we have shown that a Sox17-downstream target Rasip1 contributes to cluster formation and the hematopoietic activity of IAHC cells in the E10.5 AGM region of the midgestation mouse embryo.

## Conclusion

In summary, the *Rasip1* is identified as a Sox17 downstream gene and found to upregulate, although transient, the hematopoietic activity of HSC-containing IAHC cells of the dorsal aorta’s AGM region in midgestation (E10.5) mouse embryo.

## List of abbreviations

HSCs: hematopoietic stem cells
IAHCs: intra-aortic hematopoietic cell clusters
AGM: aorta-gonad-mesonephros
HSPCs: hematopoietic stem/progenitor cells
E: embryonic day
DMEM: Dulbecco’s modified Eagle’s medium
RT-PCR: reverse transcription-polymerase chain reaction
Tam: tamoxifen
shRNA: short hairpin RNA
Luc: luciferase
FACS: fluorescence-activated cell sorting
VE-cad: vascular-endothelial cadherin
ESAM: endothelial specific adhesion molecule
Notch1: Neurogenic locus notch homolog *protein* 1
IRES: internal ribosome entry site;
GFP: green fluorescent protein

## Declarations

### Ethics approval and consent to participate

Animal experiments were performed under institutional guidelines approved by the Animal care Committee of Tokyo Medical and Dental University (approval number: A2018-265C, A2019-108C, A2021-177C).

### Consent for publication

Not applicable.

### Availability of data and materials

RNA-sequencing data were deposited in the DNA Data Bank of Japan with the accession no. XXXXXXX (Sox17(+)_IAHCcells: Sox17 (GFP)^+^ CD45^low^c-Kit^high^ cells) and YYYYYYYY (Sox17(-)_IAHCcells: Sox17 (GFP)^−^ CD45^low^c-Kit^high^ cells). Further information and requests for resources and reagents should be directed to the authors: Ikuo Nobuhisa and Tetsuya Taga.

### Competing interests

Not applicable.

### Ethics approval and consent to participate

Animal experiments were performed under institutional guidelines approved by the Animal care Committee of Tokyo Medical and Dental University (approval number: A2018-265C, A2019-108C, A2021-177C).

### Funding

This work was supported by grants from the Ministry of Education, Culture, Sports, Science and Technology of Japan [26440118 and 18K06249 (I.N.), 22130008, 15H04292, and 18H02678 (T.T.)], Nanken-Kyoten, TMDU (Grant number, H26-A39, H27-A35, H28-A11).

### Author contributions

GM, IN, KS, RT, AI, YK, and MK conducted the in vivo and in vitro experiments. GM, IN, and TT planned the experiments. MA, MO, and AI performed RNA sequencing analysis. GM, IN, and TT wrote the manuscript. All authors read and approved the final manuscript.

## Acknowledgments

We thank Dr. T. Nakano for the OP9 cells, Ms. K. Inoue for technical assistance, and Ms. M. Makino for secretarial assistance. We also thank Ms. H. Saito to prepare Sox17^GFP/+^ mouse embryos.

## Authors’ information

The present address of Kiyoka Saito is Laboratory of Proteostasis in Stem Cell, International Research Center for Medical Sciences, Kumamoto University, 2-2-1 Honjo, Chuo-ku, Kumamoto 860-0811, Japan.

The present address of Ayumi Itabashi is the Department of Immunology, Graduate School of Medicine and Faculty of Medicine, University of Tokyo, 7-3-1, Hongo, Bunkyo-ku, Tokyo 113-0033, Japan.

The present address of Mitsujiro Osawa is Thyas Co. Ltd., Kyoto 606-8501, Japan.

---

**Supplementary Fig. 1:**
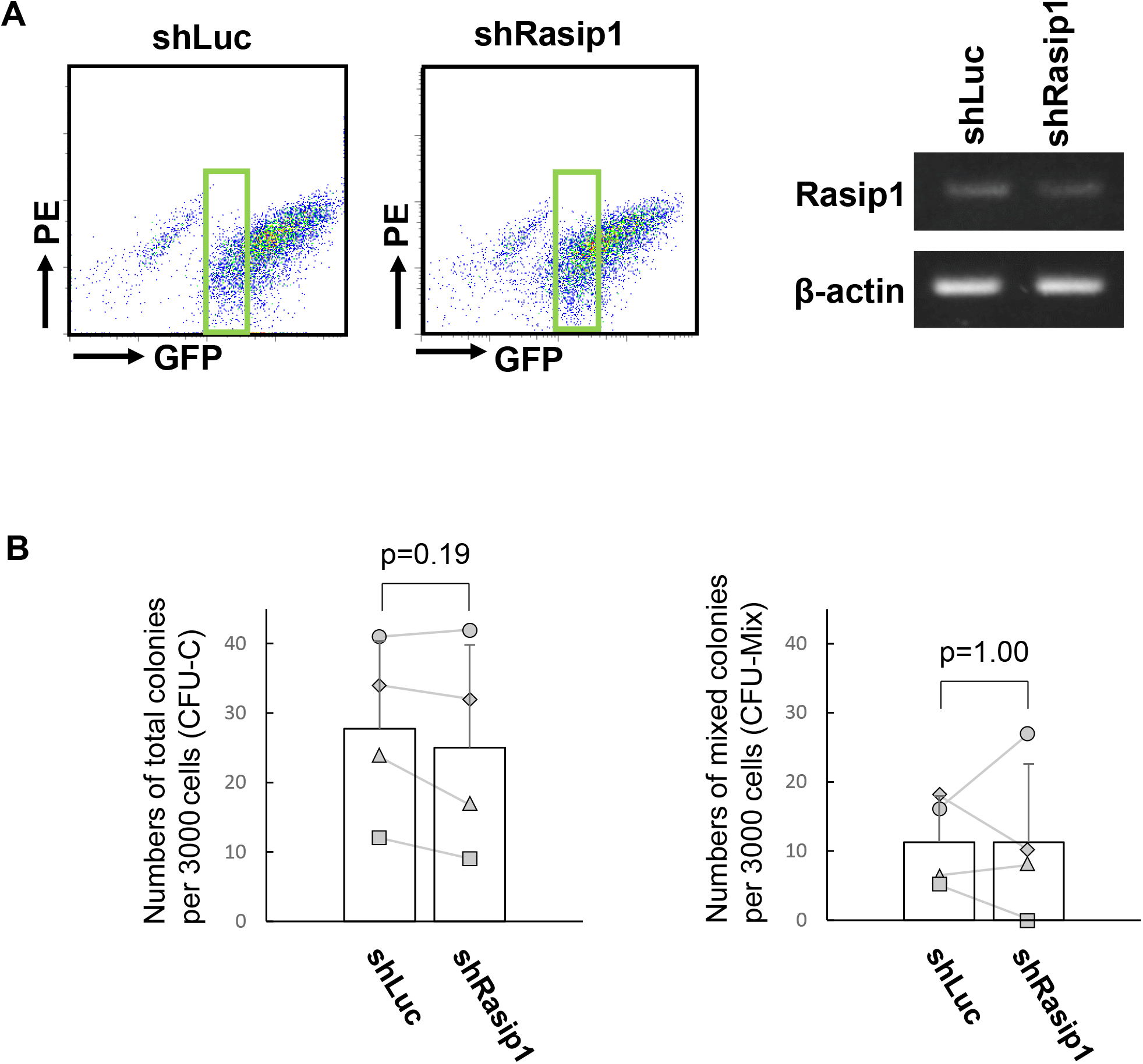
**A**. shRasip1-IRES-GFP intensity correlates with Rasip1 KD efficiency. shLuc- or shRasip1-transduced GFP^+^ cells were recovered by FACS. As indicated in the FACS profile, the light green squares show a GFP^+^ cell population with low intensity. Expression of Rasip1 in shLuc- and shRasip1-transduced cells was analyzed by RT-PCR and normalized by β-actin gene expression. RT-PCR result is in the static image that is generated by a mirror-reversal of an original across a horizontal axis, due to the sample order in the electrophoresis. **B**. Sorted GFP^+^ cells (3.0 × 10^3^) were embedded in a semi-solid medium. The number of total (CFU-C) and multilineage (CFU-Mix) colonies were scored after 7 days of culture. The pair of colony numbers (sh-Luc and shRasip1) in each experimental group were plotted and connected by a line. The different marks indicate the different individual embryos (n=4, 4 individual liters of mice from which each experimental group was set up). *p≤0.05; **p≤0.01.

**Additional file 1: Supplementary Table S1**. Table S1 shows genes expressed at high levels in Sox17 (GFP)^+^ CD45^low^c-Kit^high^ cells of E10.5 Sox17-GFP embryo as compared with Sox17 (GFP)^−^ CD45^low^c-Kit^high^ cells.

## Notes

### Competing Interest Statement

The authors have declared no competing interest.

## References

1 Dzierzak E, Speck NA. Of lineage and legacy: the development of mammalian hematopoietic stem cells. Nature Immunology. 2008;9:129–36.

2 Ferkowicz MJ, Yoder MC. Blood island formation: longstanding observations and modern interpretations. Exp. Hematol. 2005;33:1041–7.

3 de Bruijn MF, Speck NA, Peeters MC, Dzierzak E. Definitive hematopoietic stem cell first develops within the major arterial regions of the mouse embryo. EMBO J. 2000;19(11):2465–74.

4 Swiers G, Rode C, Azzoni E, de Brujin MF. A short history of hemogenic endothelium. Blood Cells Mol Dis. 2013;51(4):206–12

5 Yokomizo T, Yamada-Inagawa T, Yzaguirre AD, Chen MJ, Speck NA, Dzierzak E. Three-dimensional cartography of hematopoietic clusters in the vasculature of whole mouse embryos. Development. 2010;137(21):3651–61.

6 Bertrand JY, Giroux S, Golub R, Claine M, Boucontet L, Godin I, Cumano A. Characterization of purified intraembryonic hematopoietic stem cells as a tool to define their site of origin. Proc Natl Acad Sci USA. 2005;102(1):134–9.

7 Rybtsov S, Sobiesak M, Taoudi S, Souilhol C, Senserrich J, Liakhovitskaia A, Ivanovs A, Frampton J, Zao S, Medvinsky A. Hierarchical organization and early hematopoietic specification of the developing HSC lineage in the AGM region. J Exp Med. 2011;208(6):1305–15.

8 Mizuochi C, Fraser S, Biasch K, Horio Y, Kikushige Y, Tani K, Akashi K, Tavian M, Sugiyama D. Intra-Aortic Clusters Undergo Endothelial to Hematopoietic Phenotypic Transition during Early Embryogenesis. PLoS ONE. 2012;7(4):e35763.

9 Chen M, Yokomizo T, Ziegler B, Dzierzak E, Speck NA. Runx1 is required for the endothelial to haemapoietic cell transition but not thereafter. Nature. 2009;457(7231):887–91.

10 Nobuhisa I, Osawa M, Uemura M, Kishikawa Y, Anani M, Harada K, Takagi H, Saito K, Kanai-Azuma M, Kanai Y, Iwama A, Taga T. Sox17-mediated maintenance of intra-aortic hematopoietic cell clusters. Mol Cell Biol. 2014;34(11):1976–90.

11 Taoudi S, Gonneau C, Moore K, Sheridan JM, Blackburn CC, Taylor E, Medvinsky A. Extensive hematopoietic stem cell generation in the AGM region via maturation of VE cadherin+CD45+pre-definitive HSCs. Cell Stem Cell. 2008;3(1):99–108.

12 Taoudi S, Morrison AM, Inoue H, Gribi R, Ure J, Medvinsky A. Progressive divergence of definitive hematopoietic stem cells from the endothelial compartment does not depend on contact with the fetal liver. Development. 2005;132(18):4179–91.

13 Nobuhisa I, Otsu N, Okada S, Nakagata N, Taga T. Identification of a population of cells with hematopoietic stem cell properties in mouse aorta-gonad-mesonephros cultures. Exp Cell Res. 2007;313(5):965–74.

14 Otsu N, Nobuhisa I, Mochita M, Taga T. Inhibitory effects of homeodomain-interacting protein kinase 2 on the aorta-gonad-mesonephros hematopoiesis. Exp Cell Res. 2007;313(1):88–97.

15 Mukouyama Y, Hara T, Xu M, Tamura K, Donovan PJ, Kim H, Kogo H, Tsuji K, Nakahata T, Miyajima A. In vitro expansion of murine multipotential hematopoietic progenitors from the embryonic aorta-gonad-mesonephros region. Immunity. 1998; (1):105–14.

16 Nobuhisa I, Yamasaki S, Ramadan A, Taga T. CD45^low^c-Kit^high^ cells have hematopoietic properties in the mouse aorta-gonad mesonephros region. Exp Cell Res. 2012;318(6):705–15.

17 Saito K, Nobuhisa I, Harada K, Takahashi S, Anani M, Lickert H, Kanai-Azuma M, Kanai Y, Taga T. Maintenance of hematopoietic stem and progenitor cells in fetal intra-aortic hematopoietic clusters by Sox17-Notch1-Hes1 axis. Exp Cell Res. 2018;365(1):145–55.

18 Tam P, Kanai-Azuma M, Kanai Y. Early endoderm development in vertebrates: lineage differentiation and morphogenetic function. Curr Opin Genet Dev. 2003;13(4):393–400.

19 Yasunaga M, Tada S, Torikai-Nishikawa S, Nakano Y, Okada M, Jakt LM, Nishikawa S, Chiba T, Era T, Nishikawa S. Induction and monitoring of definitive and visceral endoderm differentiation of mouse ES cells. Nat Biotechnol. 2005;23(12):1542–50.

20 Kanai-Azuma M, Kanai Y, Gad JM, Tajima Y, Taya C, Kurohmaru M, Sanai Y, Yonehara H, Yazaki K, Tam PPL, Hayashi Y. Depletion of definitive gut endoderm in Sox17-null mutant mice. Development. 2002;129(10):2367–79.

21 Clarke R, Yzaguirre A, Yashiro-Ohtani Y, Bondue A, Blanpain C, Pear W, Speck NA, Keller G. The expression of Sox17 identifies and regulates hemogenic endothelium. Nat Cell Biol. 2013;15(5):502–10.

22 Kumano K, Chiba S, Kunisato A, Sata M, Saito T, Nakagami-Yamaguchi E, Yamaguchi T, Masuda S, Shimizu K, Takahashi T, Ogawa S, Hamada Y, Hirai H. Notch1 but not Notch2 is essential for generating hematopoietic stem cells from endothelial cells. Immunity. 2003;18(5):699–711.

23 Takahashi S, Nobuhisa I, Saito K, Melig G, Itabashi A, Harada K, Osawa M, Endo TA, Iwama A, Taga T. Sox17-mediated expression of adherent molecules is required for the maintenance of undifferentiated hematopoietic cluster formation in the midgestation mouse embryo. Differentiation. 2020;115:53–61.

24 Harada K, Nobuhisa I, Anani M, Saito K, Taga T. Thrombopoietin contributes to the formation and the maintenance of hematopoietic progenitor-containing cell clusters in the intra-gonad-mesonephros region. Cytokine. 2017;95:35–42.

25 Anani M, Nobuhisa I, Osawa M, Iwama A, Harada K, Saito K, Taga T. Sox17 as a candidate regulator of myeloid restricted differentiation potential. Dev Growth Differ. 2014;56:46979.

26 Kayamori K, Nagai Y, Zhong C, Kaito S, Shinoda D, Koide S, Kuribayashi W, Oshima M, Nakajima-Takagi Y, Yamashita M, Mimura N, Becker HJ, Izawa K, Yamazaki S, Iwano S, Miyawaki A, Ito R, Tohyama K, Lennox W, Sheedy J, Weetall M, Sakaida E, Yokote K, Iwama A. DHODH inhibition synergizes with DNA-demethylating agents in the treatment of myelodysplastic syndromes. Blood Adv. 2021;5(2):438–50.

27 Kim I, Saunders TL, Morrison SJ. Sox17 dependence distinguishes the transcriptional regulation of fetal from adult hematopoietic stem cells. Cell. 2007;130(3):470–83.

28 He S, Kim I, Lim MS, Morrison SJ. Sox17 expression confers self-renewal potential and fetal stem cell characteristics upon adult hematopoietic progenitors. Genes Dev. 2011; 25(15):1613–27.

29 Morita S, Kojima T, Kitamura T. Plat-E: an efficient and stable system for transient packaging of retroviruses. Gene Ther. 2000;7(12):1063–6.

30 Nakajima-Takagi Y, Osawa M, Oshima M, Takagi H, Miyagi S, Endoh M, Endo TA, Takayama N, Eto K, Toyoda T, Koseki H, Nakauchi H, Iwama A. Role of SOX17 in hematopoietic development from human embryonic stem cells. Blood. 2013;121(3):447–58.

31 Brummelkamp TR, Bernards R, Agami R. A system for stable expression of short interfering RNAs in mammalian cells. Science. 2002;296(5567):550–3.

32 Wu W, Bi C, Credille KM, Manro JR, Peek VL, Donoho GP, Yan L, Wijsman JA, Yan SB, Walgren RA. Inhibition of tumor growth and metastasis in non–small cell lung cancer by LY2801653, an inhibitor of several oncokinases, including MET. Clin Cancer Res. 2013;19(20):5699–710.

33 Post A, Pannekoek WJ, Ross SH, Verlaan I, Brouwer PM, Bos JL. Rasip1 mediates Rap1 regulation of Rho in endothelial barrier function through ArhGAP29. Proc Natl Acd Sci USA. 2013;110(28):11427–32.

34 Xu K, Sacharidou A, Fu S, Chong DC, Skaug B, Chen ZJ, Davis GE, Cleaver. O. Blood vessel tubulogenesis requires Rasip1 regulation of GTPase signaling. Dev Cell. 2011;19;20(4):526–39.

35 Kreuk BJ, Gingras AJ, Knight J, Liu J, Gingras AC, Ginsberg MH, Heart of glass anchors Rasip1 at endothelial cell-cell junctions to support vascular integrity. Elife 2016;5:e11394.

36 Wilson CW, Parker LH, Hall CJ, Smyczek T, Mak J, Crow A, Posthuma G, De Mazière A, Sagolla M, Chalouni C, Vitorino P, Roose-Girma M, Warming S, Klumperman J, Crosier PS, Ye W. Rasip1 regulates vertebrate vascular endothelial junction stability through Epac1-Rap1 signaling. Blood. 2013;122(22):3678–90.

37 Koo Y, Barry DM, Xu K, Tanigaki K, Davis GE, Mineo C, Cleaver O. Rasip1 is essential to blood vessel stability and angiogenic blood vessel growth. Angiogenesis. 2016 (2):173–90.

38 Xu K, Chong DC, Rankin SA, Zorn AM, Cleaver O. Rasip1 is required for endothelial cell motility, angiogenesis, and vessel formation. Dev Biol. 2009;329(2): 269–79.

39 Mitin NY, Ramocki MB, Zullo AJ, Der CJ, Konieczny SF, Taparowsky EJ. Identification and Characterization of Rain, a Novel Ras-interacting Protein with a Unique Subcellular Localization. J Biol Chem. 2004;279(21):22353–6

